# Mechanisms of surface antigenic variation in the human pathogenic fungus *Pneumocystis jirovecii*

**DOI:** 10.1101/158881

**Authors:** Emanuel Schmid-Siegert, Sophie Richard, Amanda Luraschi, Konrad Mühlethaler, Marco Pagni, Philippe M. Hauser

**Affiliations:** Vital-IT Group, SIB Swiss Institute of Bioinformatics, Lausanne, Switzerland; Institute of Microbiology, Lausanne University Hospital, Lausanne, Switzerland; Institut für Infektionskrankheiten, Universität Bern, Bern, Switzerland

**Keywords:** *Pneumocystis carinii*, major surface glycoprotein, adhesin, subtelomere, PCP, mosaicism, gene conversion, gene exchange, telomere exchange, PacBio sequencing

## Abstract

Microbial pathogens commonly escape the human immune system by varying surface proteins. We investigated the mechanisms used for that purpose by *Pneumocystis jirovecii.* This uncultivable fungus is an obligate pulmonary pathogen which causes pneumonia in immuno-compromised individuals, a major life-threatening infection. Long-read PacBio sequencing was used to assemble a core of subtelomeres of a single *P. jirovecii* strain from a bronchoalveolar lavage fluid specimen of a single patient. A total of 113 genes encoding surface proteins were identified, including 28 pseudogenes. These genes formed a subtelomeric gene superfamily which included five families encoding adhesive GPI-anchored glycoproteins, and one family encoding excreted glycoproteins. Numerical analyses suggested that diversification of the glycoproteins relies on mosaic genes created by ectopic recombination, and occurs only within each family. DNA motifs suggested that all genes are expressed independently, except those of the family encoding the most abundant surface glycoproteins which are subject to mutually exclusive expression. PCR analyses showed that exchange of the expressed gene of the latter family occurs frequently, possibly favoured by the location of the genes proximal to the telomere because this allows concomitant telomere exchange. Our observations suggest that (i) *P. jirovecii* cell surface is made of a complex mixture of different surface proteins, with a majority of a single isoform of the most abundant glycoprotein, (ii) genetic mosaicism within each family ensures variation of the glycoproteins, and (iii) the strategy of the fungus consists in the continuous production of new subpopulations composed of cells which are antigenically different.

## Importance

*Pneumocystis jirovecii* is a fungus causing severe pneumonia in immuno-compromised individuals. It is the second most frequent life-threatening invasive fungal infection. We have studied the mechanisms of antigenic variation used by this pathogen to escape the human immune system, a strategy commonly used by pathogenic microorganisms. Using a new DNA sequencing technology generating long reads, we could characterize the highly repetitive gene families encoding the proteins which are present on the cellular surface of this pest. These gene families are localized in the regions close to the ends of all chromosomes, the subtelomeres. Such chromosomal localization was found to favour genetic recombinations between members of each gene family and allow diversification of these proteins continuously overtime. This pathogen proves to use a strategy of antigenic variation consisting in the continuous production of new subpopulations composed of cells which are antigenically different. Such strategy is unique among human pathogens.

## Introduction

*Pneumocystis jirovecii* is a fungus colonizing specifically human lungs. It has developed strategies to survive in healthy human lungs, at least transiently, and can turn into a deadly pathogen causing pneumonia in individuals with debilitated immune system (1-4). This disease is the second most frequent life-threatening invasive fungal infection with ca. 400’000 cases per year worldwide (5). However, the biology of this pest remains difficult to study in the lab because of the lack of any established methods for *in vitro* culture. Recent progresses in understanding *P. jirovecii* biology strongly benefitted from the publication of two assemblies of its genome from two different clinical samples (4, 6).

In contrast to other pathogenic fungi, the cells of *P. jirovecii* lack chitin as well as glucans during part of the cell cycle, which may avoid eliciting innate and acquired immune responses (4). Moreover, a mechanism of surface antigenic variation, to which ca 5% of the genome is dedicated, seems crucial to escape from the human immune system during colonization, although it has not been understood in details so far. Surface antigenic variation is a common strategy among major microbial human pathogens, for example in *Plasmodium*, *Trypanosoma*, *Candida*, *Neisseria*, and *Borrelia*. It relies on various genetic and/or epigenetic mechanisms aimed at expressing only one or few of them at once (7). Such systems often involve gene families encoding surface antigens localized at subtelomeres, presumably because these regions of the genome are prone to gene silencing, which is used for mutually exclusive expression, and possibly to enhanced mutagenesis (8). Moreover, the formation of clusters of telomeres at the nuclear periphery may favour ectopic recombinations (8), which can be responsible for the generation of new mosaic antigens.

Surface antigenic variation has been previously studied on a limited set of genes in *Pneumocystis carinii* infecting specifically rats. The molecular mechanism was then assumed to be also active in *P. jirovecii*, as suggested by studies using PCR-based technologies. Antigen diversity was believed to be generated by recombination between members of a single family of ca. 80 subtelomeric genes encoding isoforms of the major surface glycoprotein (*msg*) (9-11). A single of these isoforms would be expressed in each cell thanks to its localization downstream of a subtelomeric expression site, the upstream conserved element (UCS) present at a single copy in the genome. The UCS includes the promoter of transcription, the protein start, and the leader sequence responsible for translocation of the protein into the endoplasmic reticulum for final incorporation into the cell wall (12, 13). The mechanism for exchange of the expressed *msg* gene is thought to be by recombination at a 33 bps long sequence which is present both at the end of UCS and beginning of each *msg* (conserved recombination junction element, CRJE). The exchange of the expressed gene seems relatively frequent and would explain how different *msg* genes can be expressed in each population (12). The CRJE sequence encodes at its end a potential lysine-arginine recognition site for Kexin endonuclease which might be involved in the maturation of the antigen. Kutty *et al* (14) provided evidence for frequent recombinations among *msg* genes creating potentially mosaic genes. All these observations were made using conventional cloning procedures and PCRs, and these mechanisms have yet to be understood in a more extensive genomic context.

The first genome sequence of *P. jirovecii* released was obtained using technologies generating short reads which prevented assembly of long repetitive sequences such as centromeres, telomeres, and subtelomeres including *msg* genes (6). A second study used a mixture of techniques which generated nearly complete chromosomes of *P. jirovecii*, *P. carinii*, and *Pneumocystis murina* (infecting specifically mice) (4). These latter authors used PCRs to reconstruct the subtelomeres which allowed discovering new subtelomeric gene families related to *msg*. However, they did not investigate the function of these proteins, the mechanisms involved in their expression and gene variation, or the global strategy of antigenic variation of these fungi.

The aim of the present study was to analyse in details the mechanisms of surface antigenic variation in *P. jirovecii*. To that purpose, we used the PacBio sequencing technology generating long DNA reads to assemble a set of subtelomeres of a single *P. jirovecii* strain from a bronchoalveolar lavage fluid specimen (BALF) of a single patient. The analysis of this dataset and laboratory experiments permit a new classification and the characterization of six subtelomeric *msg* families, demonstrate the presence of pseudogenes, and provide important new insights into the molecular mechanisms responsible for antigenic variation. Moreover, our observations suggest a unique strategy of antigenic variation consisting in the continuous production of new subpopulations composed of cells which are antigenically different. This strategy may be associated to the particular non-sterile niche within lungs.

## Results

Most if not all *P. jirovecii* infections are polyclonal (15). In order to facilitate the study of the mechanisms of antigenic variation, one patient infected with a vastly dominant strain was selected by multitarget genotyping. The genome of a single *P. jirovecii* strain was assembled into 219 contigs using PacBio sequencing and a dedicated bioinformatics strategy for reads processing.

### Identification of subtelomeric *msg* genes and pseudogenes

Automated gene prediction performed poorly in the subtelomeric regions as compared to the core of the genome, due to abundant stretches of low-complexity DNA, numerous pseudogenes, residual assembly errors in homopolymers, and the lack of a start codon in many *msg* genes. The *msg* genes were detected by sequence homology using generalized profiles (16) derived from previously published sequences. A total of 113 *msg* genes with sizes ranging from 331 to 3337 bps were found on 37 different contigs, only two genes being perfectly identical (*msg*52 and 61; Additional file 1: Table S1). Most of them (N=85) contained a single large exon and zero to two small exons at their 5’ end. The remaining 28 genes harboured many stop codons in all frames and were considered as pseudogenes (Additional file 2: supplementary note 1).

### Characterization of the *msg* gene families

We are proposing a classification of the *msg* genes into six families (Table 1) based on of the integration of four independent lines of evidence: sequence homology, gene structure, protein property, and recombination events. The global picture that emerged is coherent and the details on the different points are presented below.

**Table 1.** Characteristics of the *msg* families identified in *P. jirovecii.*

| Family name | Color in figures | Gene |  |  |  |  |  |  | Protein |  |  |  |  |  |  |
| --- | --- | --- | --- | --- | --- | --- | --- | --- | --- | --- | --- | --- | --- | --- | --- |
| | | No. genes full-length / partial / pseudo- | Mean full-length (bps) ± st dev | Location in subtelomere relatively to telomere | Presumptive TATA box (bps to ATG, range) | CRJE | No. 5'-end introns | Average pairwise identity (%) ± st dev | Signal peptide | C-terminus | | | GPI-anchor signal | No. N-glycosylation site | Average pairwise identity (%) ± st dev |
|  |  |  |  |  |  |  |  |  |  | ST-rich region | ST-rich region | PE-rich region |  |  |  |
| *msg*-I | | 11 / 16 / 16 | 3071 ± 39 | proximal | - <sup>a</sup> | + | 0 | 71 ± 7 | - <sup>b</sup> | + | + | - | + | 4-10 | 54 ± 8 |
| *msg*-II | | 11 / 3 / 4 | 3155 ± 31 <sup>b</sup> | central | 21-28 | - | 2 | 83 ± 13 | + | + | + | - | + | 2-14 | 73 ± 16 |
| *msg*-III | | 7 / 2 / 1 | 3146 ± 55 | central | 18-24 | - | 2 | 83 ± 10 | + | + | + | - | + | 7-11 | 70 ± 13 |
| *msg*-IV | | 6 / 1 / 2 | 2023 ± 45 | central | 29-36 | - | 1 | 72 ± 14 | + | - | - | - | - | 0-8 | 49 ± 17 |
| *msg*-V | | 8 / 6 / 1 | 3056 ± 126 | central | 30-67 | - | 1 | 66 ± 5 | + | - | + | + | + | 5-12 | 44 ± 4 |
| *msg*-VI | | 6 / 1 / 0 | 1222 ± 189 | distal | 33-56 | - | 1 | 45 ± 7 | + | - | + | + | + | 0-1 | 21 ± 5 |
| *msg* outlier |  | 6 / 1 / 4 | variable | central/distal | NA <sup>c</sup> | +/- | variable | NA | +/- | +/- | +/- | - | +/- | variable | NA |

<sup>a</sup> The promoter including the signal peptide for family I is within the UCS present at a single copy per genome.
<sup>b</sup> The *msg3* gene was not used to calculate this value because it is ca. 900 pbs shorter than the other genes of the family, although it presents all features of the family (see alignment in Figure S2).
<sup>c</sup> Not applicable.

Figure 1a shows the results of the analysis of 61 *msg* genes containing an exon equal or larger than 1.6 kb. Based on the multiple sequence alignments (MSAs) of the CDS and of their predicted proteins, two phylogenetic trees were computed using RAxML. The different gene families are clearly individualised as clades, with the exceptions of (i) *msg*-II which appears as a sub-clade of *msg*-I, and (ii) *msg*-I which seems to include two sub-clades. Using an alternative classification method that does not rely on a single particular MSA (JACOP, Fig. 1a), the placement of *msg*-II as a sub-clade of *msg*-I was not confirmed, whereas the sub-clades of *msg*-I were. Owing on the differences in the gene structures and on the recombination events reported below, we believe that (i) *msg*-I and *msg*–II should be treated separately, and (ii) *msg*-I should be considered as a single family including two sub-clades. Figure 1b shows the analysis of trimmed CDS sequences allowing the placement of the *msg*-VI family which appeared as a clade on its own, while the classification of the other families remained essentially unchanged. Figure S1 shows that most pseudogenes could be attributed to one *msg* family and their often longer branches further account for their pseudogenic nature (Additional file 2).

**Fig. 1.**
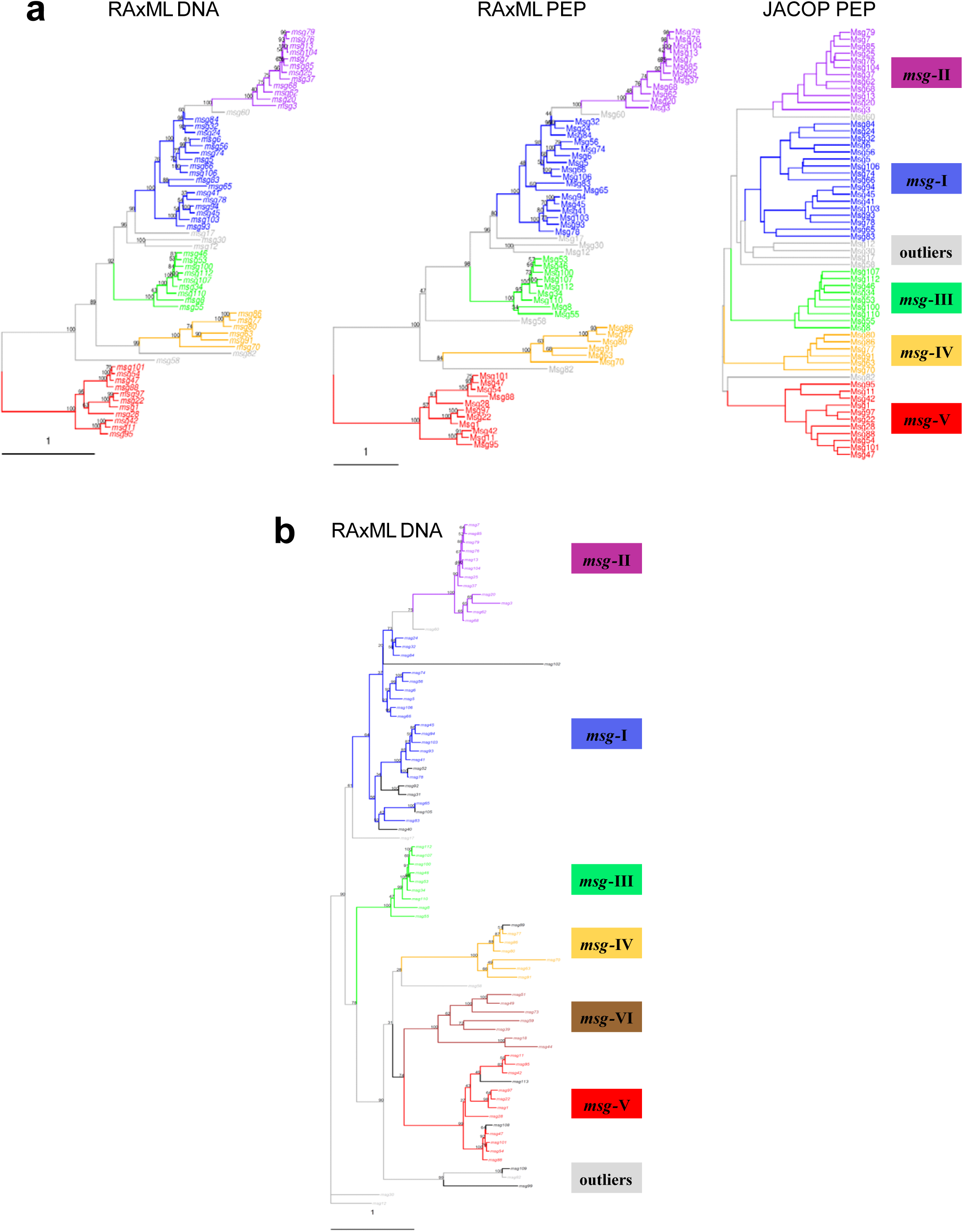
Classification trees of *P. jirovecii msg* genes and Msg proteins. The different families are represented in colours and their characteristics are summarised in Table 1. A few unclassified outliers are in grey. Scale, mean substitution / site. (**a**) RAxML DNA and PEP are maximum likelihood trees of nucleotide and amino acid sequences of the 61 genes with an exon larger than 1.6 kb. Members of family V were defined as the out-group (1000 bootstraps). JACOP PEP is a hierarchical classification based on local sequence similarity, a method that does not rely on a particular multiple sequence alignment. (**b**) Maximum likelihood tree of the 61 genes with an exon larger than 1.6 kb, plus 18 genes with an exon smaller than 1.6 kb. The sequences were trimmed from position 1540 of the first alignment up to their end, and re-aligned to construct the tree (1000 bootstraps). Seven of the 18 genes with an exon smaller than 1.6 kb constitute the *msg* family VI shown in brown, whereas the remaining 11 shown in black belong to the other *msg* families.

Manual curation of the *msg* genes led to their classification in full-length, partial, and pseudogenes (Table S1). Table 1 shows the characteristics of each family identified by the analysis of the sequences of the full-length genes, as well as of their alignments (Fig. S2). Except those of the family *msg-*I, each *msg* gene presented one or two introns at its 5’ end, as well as a presumptive TATA box upstream of the ATG and an initiator motif (Cap signal) at presumptive sites of initiation transcription (Fig. 2a and S2). The members of the family I had only the CRJE (conserved recombination junction element) at the beginning of their single exon. These observations suggested that members of family I can be expressed only upon recombination of their CRJE with that of the single copy UCS which encompasses a promoter, whereas all members of the other five families are expressed independently. Three of the six full-length outlier genes seemed not expressed since they had no CRJE and missed a TATA box (Table S1). Twenty-six partial genes were truncated by the end of the contig so that only three *bona fide* partial genes were identified, which, however, missed TATA box, signal peptide, and/or GPI-anchor signal, and thus were probably not expressed or not correctly processed (*msg*44, 89, and 99).

**Fig. 2.**
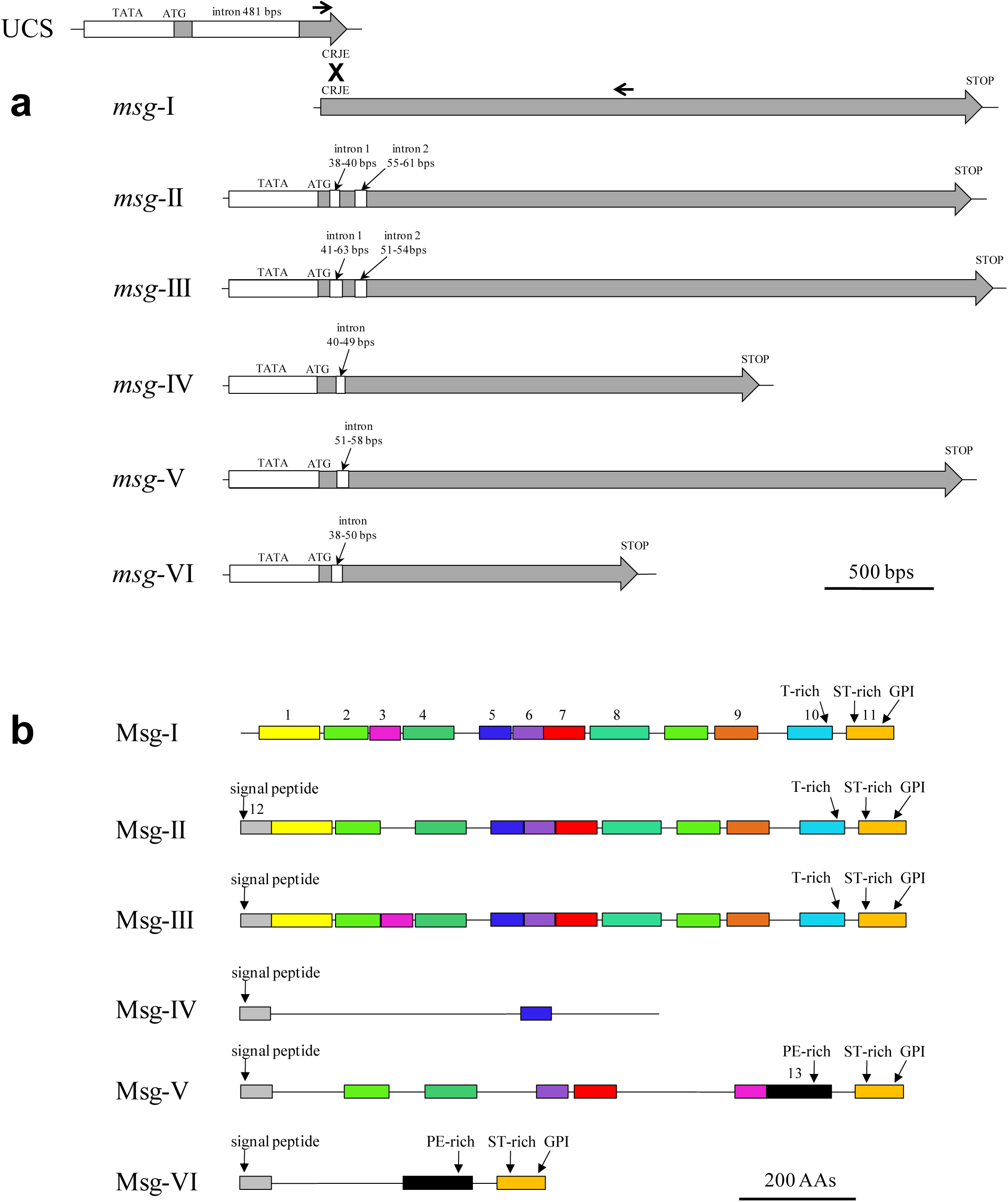
Diagrams of the structure of *P. jirovecii msg* genes and Msg proteins belonging to families I to VI. (**a**) Features of the *msg* genes of each family derived from the analysis of the full-length genes. The UCS and recombination between CRJE sequences are figured for family I. The approximate position of PCR primers used for identification of the *msg*-I expressed genes linked to the UCS are shown by arrows (Supplementary note 4). (**b**) Features of Msg proteins of each family derived from the analyses of the full-length proteins. The 13 domains identified using MEME analysis are shown. The logos of these domains are shown in Figure S4.

### Characterization of the Msg protein families

Analysis of the sequences and alignments (Fig. S3) of the full-length proteins of each family revealed that each Msg protein, except those of family I, presented a signal peptide at its N-terminus (Fig. 2b). Proteins of family I probably acquire a signal peptide upon fusion of their encoding gene with the UCS. Except those of family IV, each Msg protein presented a GPI-anchor signal at its C-terminus. These observations suggested that all Msg proteins are attached externally to the cell wall, except those of family IV which would be secreted in the environment or attached to the cell wall through another mechanism than GPI.

The possible conservation of motifs among the proteins of the six families was investigated using Multiple Expectation–Maximization for Motif Elicitation (MEME analysis) (17). Thirteen conserved motifs were identified which arrangement was fairly diagnostic within each family (Fig. 2b). Most motifs included several conserved cysteines and leucines, which resembled to the previously identified Pfam MSG domain (Fig. S4). Interestingly, conserved leucines were often separated by two to six residues. The beginning of motif 10 corresponded to the end of the previously identified Pfam Msg2_C domain. Accordingly, Pfam predictions identified one to five MSG domains, often partial, per protein of all families, and a single Msg2_C domain in each Msg-I protein (Fig. S5; Table S2). The Msg2_C domain was not predicted in families II and III although they harboured the corresponding motif 10, suggesting that this domain is divergent in these families. Ncoils predictor revealed three to five coiled-coil motifs spread along members of families I, II, and III, whereas unstructured regions were predicted at the C-terminus of Msg proteins of families I, III, V, and VI (Fig. S5).

Except those of family IV, each Msg protein harboured at its C-terminus two MEME motifs which included a region enriched in specific residues: threonine (T-rich; motif 10), serine and threonine (ST-rich; 11), or proline and glutamine (PE-rich; 13)(Fig. 2b; Table 1). The T-rich region in family I included generally a stretch of nine to 15 Ts, which was not present in families II and III (Fig. S3). The PE-rich region in family V was enriched in proline residues relatively to that present in family VI (Fig. S3). Four to 14 potential sites of nitrogen-linked glycosylation of asparagines were predicted to be present in each Msg protein, except in family VI which presented no or only one such site (Table 1; Fig. S3). The localization of these glycosylation sites was widespread along the protein and fairly conserved within each family (Fig. S3).

### Arrangement of the *msg* families within the subtelomeres

Consistent with a subtelomeric localization, the *msg* genes were grouped at one end of their contig when flanking non-*msg* genes were also present (in 20 of 37 contigs; Fig. 3 and S6a). All *msg* genes identified were oriented towards one end of the contig, *i.e.* presumably towards the telomere (no telomeric repeats were identified for an unknown reason; Supplementary note 2). Except pseudogenes which were dispersed all over the subtelomeres, all members of family I were the closest to the end of their contig, *i.e.* proximal to the telomere (Fig. 3 and S6). By contrast, all members of family VI were the closest to the flanking non-*msg* genes present on their contig, *i.e.* distal to the telomere. Members of the four remaining families were localized centrally in the subtelomeres, between those of families I and VI. There were up to three *msg*-I genes grouped at the end of 19 contigs. Members of the other five families did not show any clear grouping patterns.

**Fig. 3.**
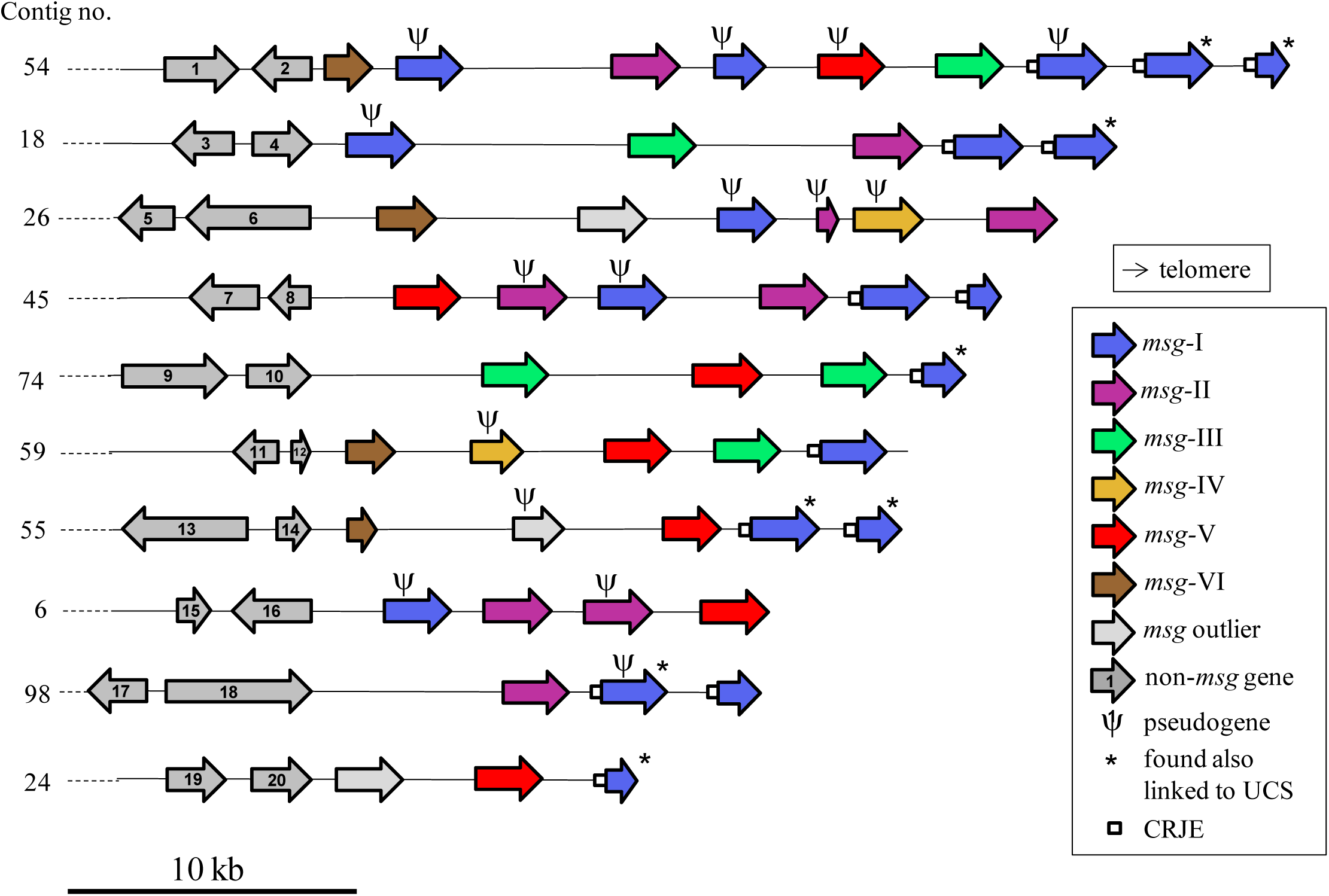
Diagrams of ten representative *P. jirovecii* assembled subtelomeres. The other 27 assembled subtelomeres are shown in Figure S6. The attribution of the contigs to the chromosomes previously described using flanking non-msg genes is given in Table S3.

### Identification of the expression site of *msg-* I genes and of the genes linked to it

Each infection by *P. jirovecii* is believed to involve a mixture of cells expressing different *msg*-I genes under the control of the expression site, *i.e.* the UCS which is present at a single copy per genome (12). Consequently, the UCS was expected to be linked to different *msg*-I genes in our DNA sample and thus to cannot be unequivocally assembled, which plausibly explains its absence from the PacBio assembly. A single UCS was retrieved from our DNA sample using PCRs based on published sequences and it could be linked to one of the PacBio contigs (Supplementary note 3). The UCS retrieved from our sample was identical to that of Ma *et al* (4), except few small changes not modifying the encoded protein (Fig. S7). Interestingly, the CRJE sequence at the end of the UCS and beginning of each *msg*-I gene presented an imperfect inverted repeat which was never pointed out so far (Fig. S7).

In order to identify the *msg*-I genes linked to the UCS in our sample, we amplified by PCR the junction between these elements using one primer within the UCS and either one primer generic for many *msg*-I genes (12), or one primer specific to a given *msg*-I gene of the PacBio assembly (Supplementary note 4; Fig. 2a). Eighteen different *msg*-I genes were found fused in frame to the UCS at the CRJE sequence, two being pseudogenes of the family I with an upstream CRJE sequence, and four new *msg*-I sequences not present in the PacBio assembly. The 12 *msg*-I genes found linked to the UCS which were present in the PacBio assembly are identified in Figures 3 and S6 by asterisks. Three specific *msg*-I genes linked to the UCS represented 74% of the subclones of the generic PCR analyzed, suggesting that sub-populations of cells expressing given *msg*-I genes were of different sizes in our sample (Supplementary note 4). These observations suggested that recombination between the CRJE sequence of the UCS with that of different *msg*-I genes occurred at a high frequency in the single *P. jirovecii* population studied here.

### Set of assembled subtelomeres

The flanking non-*msg* genes allowed attributing 20 of our 37 contigs (Fig. 3 and S6a) to 15 of the 20 full-length chromosomes described by Ma *et al* (4) because they were also present in the latter assembly (Table S3). All the remaining 17 contigs without flanking non-*msg* genes (Fig. S6b) could have been assembled from the same subtelomeres as other contigs. Thus, we assembled at least 20 subtelomeres out of the 40 potentially present in each cell. Given the presence of a large number of subpopulations expressing different *msg*-I genes in our sample, the set of subtelomeres present in each cell varied considerably. It is likely that the set we assembled corresponded to a core of subtelomeres which was present in a majority of cells of the population so that it could be assembled unequivocally.

### Recombination between *msg* genes

Evidence of recombination events between *msg*-I genes was previously provided (14). We investigated this issue among the different *msg* families using three different numerical methods: two allowing analyses of large sets of genes for screening, and one analyzing only four genes at a time for more sensitive analysis. Two to 18 potential mosaic genes and their putative parent genes were detected within each family I to IV, involving sometimes partial or pseudogenes (Fig. 4; Table 2). On the other hand, only one potential mosaic gene was identified in family V and none in family VI (P = 0.06). Eight of the 30 mosaic genes detected shared with one parent a perfectly or almost perfectly identical fragment of ca. 100 to 1000 bps, often close to the site of the predicted recombination events (Fig. 4b and S8). These latter cases suggested very recent recombination events. The putative parent genes of mosaic genes were randomly distributed among the two sub-clades of family I, suggesting that this family must be considered as a single entity (Supplementary note 5).

**Table 2.** Potential mosaic genes detected within each *msg* family ^a^.

| *msg* family | No. *msg* genes |  |  |  |  | No. potential *msg* mosaic genes |  |  |  |  |
| --- | --- | --- | --- | --- | --- | --- | --- | --- | --- | --- |
|  | Full-length | Partial | Pseudo | Total | Non-mosaic <sup>b</sup> | Full-length <sup>c</sup> | Partial | Pseudo <sup>d</sup> | Total <sup>b</sup> | % mosaic |
| I | 11 | 16 | 16 | 43 | 25 | 8 | 1 | 9 | 18 | 42 |
| II | 11 | 3 | 4 | 18 | 13 | 4 | 1 | 0 | 5 | 28 |
| III | 7 | 2 | 1 | 10 | 6 | 3 | 0 | 1 | 4 | 40 |
| IV | 6 | 1 | 2 | 9 | 7 | 1 | 0 | 1 | 2 | 22 |
| V <sup>e</sup> | 8 | 6 | 1 | 15 | 14 | 1 | 0 | 0 | 1 | 7 |
| VI <sup>e</sup> | 6 | 1 | 0 | 7 | 7 | 0 | 0 | 0 | 0 | 0 |

<sup>a</sup> Detected using the Recombination Analysis Tool and / or Bellerophon bioinformatics screening methods among three different sets of genes of each *msg* family: full-length, full-length plus partial genes, full-length plus pseudogenes.
<sup>b</sup> The number of potential mosaic genes among the *msg* families was almost significantly different ( $P = 0.06$ , Chi-square test).
<sup>c</sup> Six full-length mosaic genes were detected twice but with different pairs of putative full-length parent genes according to the set of genes analysed (four, one, one of respectively family I, II, III). One mosaic gene of family I was detected twice: once with one full-length gene and one pseudogene as parents, and once with two partial genes as parents. All ten remaining were detected only once with a pair of full-length parents.
<sup>d</sup> Six mosaic pseudogenes of family I had two pseudogenes as parents. Two of family I had one full-length gene and one pseudogene as parents. The three remaining had a pair of full-length parents.
<sup>e</sup> Several potential recombination events were detected for these two families using the more sensitive method TOPALi based on the Hidden Markov Model (Fig. S9).

**Fig. 4.**
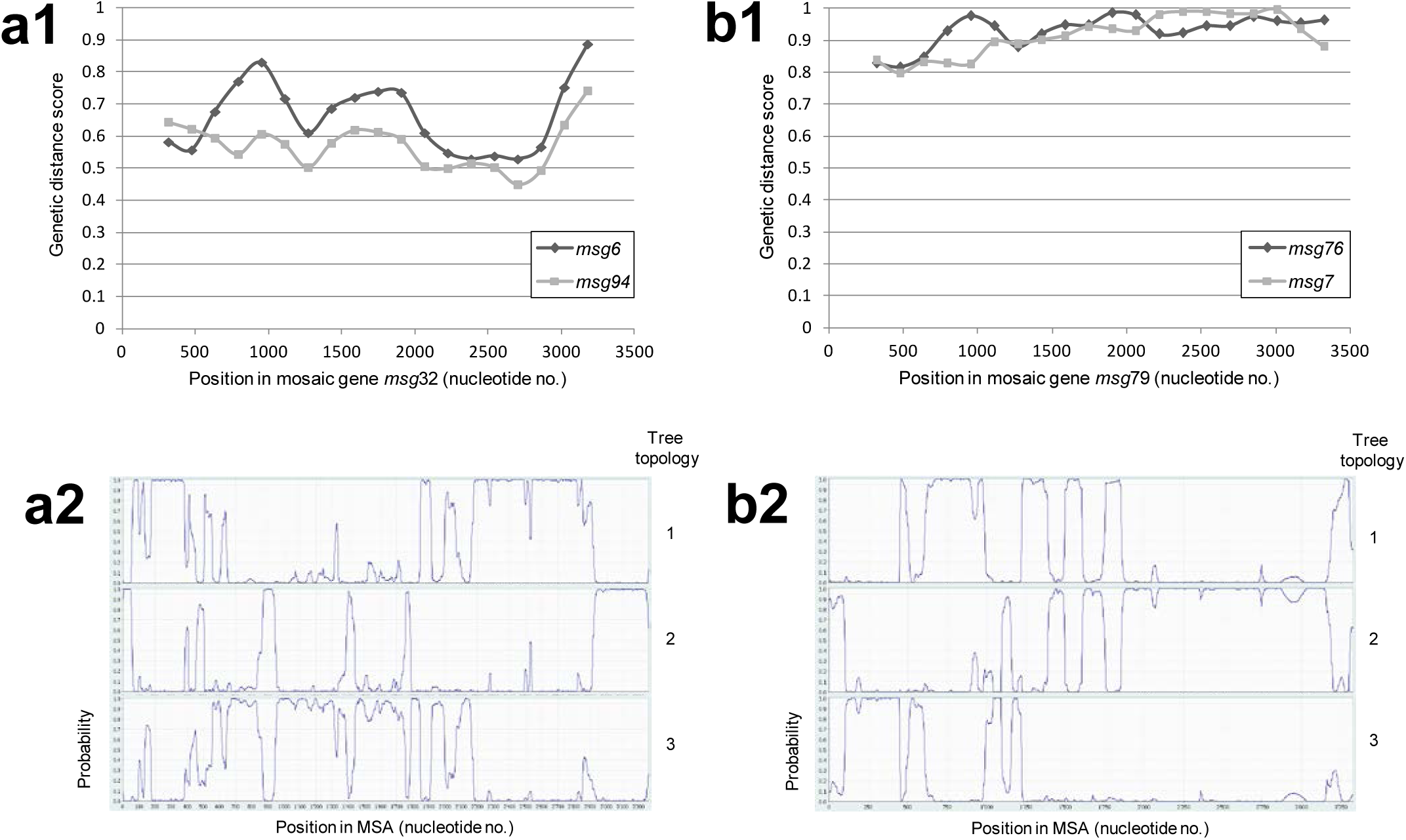
Examples of detection of potential mosaic genes. (**a**) Mosaic gene *msg*32. (**a1**) The set of 11 full-length *msg*-I genes was analyzed using the Recombination Analysis Tool. This method measures genetic distances in windows sliding along the MSA. The genetic distance scores of the putative parent genes at the middle of each window are plotted against the position in the mosaic gene. The predicted recombination site is at position ca. 600, at the cross-over of the curves. The second screening method Bellerophon, which is based on a similar analysis, identified a recombination event at position 392. (**a2**) Analysis of the mosaic gene *msg*32 with its putative parent genes together with the randomly chosen *msg*84 of the same family using the more sensitive method TOPALi based on the Hidden Markov Model. This method analyses only four sequences at a time and calculates the probabilities of the three possible tree topologies at each residue of the MSA. A recombination event is also detected at position ca. 400-600, but many other recombination events are predicted. (**b**) Mosaic gene *msg*79. This gene shares an almost identical fragment of 947 bps with its putative parent *msg*7 (see alignment in Fig. S8c). (**b1**) The set of 11 full-length *msg*-II genes was analyzed using the Recombination Analysis Tool. The predicted recombination sites are at positions ca. 400, 1300, 2100, and 3100. The Bellerophon method did not identify this mosaic gene. (**b2**) Analysis of the mosaic gene *msg*79 with its putative parent genes together with the randomly chosen *msg*85 of the same family using TOPALi based on the Hidden Markov Model. Recombination events are also detected at positions ca. 400, 1500, and 3100, but not at 2100, and other recombination events are predicted.

One to four potential recombination events per mosaic gene were generally identified using the two screening methods. These events were most often confirmed by the more sensitive method which, however, detected many other potential recombination events (Fig. 4 and S8). Consistent with the single mosaic gene detected in families V and VI, the frequency of recombination events appeared lower in these families than in the others (Fig. S9). This correlated with an average pairwise identity lower within each of these two families than within the others (45-66 versus 71-83%, Table 1). The predicted sites of the recombinations reported by all three methods were distributed randomly along the *msg* genes for all families, and did not contain any specific DNA sequence motifs (Fig. 4, S8, and S9). This suggested homologous rather than site-specific recombination events.

In contrast, we were unable to detect recombination events between different *msg* families, even using the more sensitive method (Fig. S10).

### Comparison to the *msg* superfamily previously proposed

The 146 *P. jirovecii msg* genes larger than 1.6kb reported by Ma *et al* (4), out of a total of 179, were added into our DNA phylogenetic tree. They all clustered within our families, except 11 outliers clustering with our outliers (Fig. S11). The correspondence between the two sets of families and the comparison of the two studies are detailed in the Supplementary note 6.

## Discussion

Antigenic surface variation plays a crucial role in escaping the human immune system and adhering to host cells for important microbial pathogens. In the present study, we investigated the mechanisms used by the fungus *P. jirovecii* for this purpose. Our observations show that its surface glycoproteins diversified during the evolution into a superfamily including six families each with its own structure, function, independent mosaicism, and expression mode.

### Structure and function of Msg glycoproteins

Members of the Msg family I were previously demonstrated to adhere human epithelial cell through binding to fibronectin and vitronectin (18, 19). The ST-rich regions present in *P. jirovecii* Msg glycoproteins except those of family IV are sites of oxygen-linked glycosylation commonly involved in cell to cell adhesion (20). Moreover, most of these glycoproteins were predicted to be adhesins (Supplementary note 7). Consistently, their structure fits the model of modular organization of fungal adhesins with ST-rich regions at the C-terminus and a ligand binding domain at the N-terminus (20, 21). Linder and Gustafsson (21) proposed that, in addition to their role in adhesion, the oxygen-linked glycosylations of the ST-rich region confer rigidity to the protein in order to present outward the ligand domain. Thus, the N-terminus regions of the *P. jirovecii* adhesins may correspond to ligand binding domains. The fate and function of the glycoproteins of family IV remain enigmatic since they lack the ST-rich region, are only weakly predicted as adhesins (Supplementary note 7), and may not be attached to the cell wall in absence of a GPI anchor signal. The conserved leucines separated by two to six residues present in all *msg* families are similar to leucine zipper motifs which are often involved in protein-protein non-specific binding and protein dimerization (22). This latter function is also carried out by the PE-rich region present in *msg* family V and VI (23). The conserved coiled-coil domains discovered in Msg families I to III are often involved in the formation heteromultimers and protein complexes (24, 25). On the other hand, the unstructured regions at the C-terminus present in four Msg families are not informative because these regions can have several different functions (26). These observations suggest that the Msg adhesins may form homo- or hetero-oligomers at the cell surface, possibly implying a further level of antigen variation which has never been envisaged so far.

### Mosaicism of *msg* genes

Our observations suggest that a continuous and random creation of mosaic genes by homologous recombinations occurs mostly, if not exclusively, within each *msg* family. Very interestingly within the scope of protein annotation, this mechanism permits by itself to define the members of a protein family without having to rely upon the cutting of a phylogenetic tree at an arbitrary height. The frequency of these recombinations remains to be quantified precisely, but is likely to be reduced in *msg* families V and VI. The genetic mechanisms involved in the creation of mosaic genes may include a single homologous recombination leading to a telomere exchange, or two homologous recombinations leading to a gene fragment conversion or exchange (models are shown in Fig. S12). Such recombinations could also produce partial genes if they occur between homologous regions which are not located at the same position along the recombining genes. Our results suggest that this is rare because we identified only three partial *msg* genes out of 113. This conclusion is also consistent with the fact that different motifs are conserved along the sequence of the Msg proteins of each family. Our data suggest that pseudogenes might also be involved in the generation of mosaic genes, and thus might constitute a reservoir of sequences that can be integrated into functional antigens. The pseudogenes may result from accumulation of mutations in absence of expression and thus of selective pressure. This phenomenon could be enhanced by mutation and recombination rates within the subtelomeric gene families higher than in the rest of the genome (8). The presence of the pseudogenes in the subtelomeres might simply correspond to the state between their birth and their future decay. However, they could also be maintained within the subtelomeres through indirect selective pressure because of their role as reservoir of fragments for the creation of mosaic genes.

### Mutually exclusive expression of *msg*-I genes

Our conclusions concerning the mutually exclusive expression of the *msg*-I genes are in agreement with previous studies, but bring support for the involvement of telomere exchange which has been previously hypothesized (27). The exchange of the single expressed gene by recombination at the CRJE sequences might be facilitated by the localization of the *msg*-I genes closest to the telomeres, because this may in turn facilitate telomere exchanges (a model is shown in Fig. 5). These recombinations could be homologous in nature because the full identity over 33 bps might be sufficient as it is the case in fungal cousins (28). However, they could also be site-specific because the imperfect inverted repeat present in the CRJE is a common motif used by site-specific recombinases (29). Up to three *msg*-I genes were present at the end of the subtelomeres. There is no reasons to exclude that transfer of more than one *msg*-I gene to the expression site at once also occurs, followed by polycistronic expression. The polypeptide produced could be then chopped by the endoprotease Kex1 at the end of each CRJE and each Msg-I anchored to the cell wall separately through its own GPI signal. Interestingly, we detected *msg*-I pseudogenes linked to the UCS using PCR in our sample. The cells expressing such truncated antigens may not be selected over time during the infection because of their likely deficiency in adhesion to host cells. They might constitute a cost inherent to such system of antigenic variation based on frequent recombination events.

**Fig. 5.**
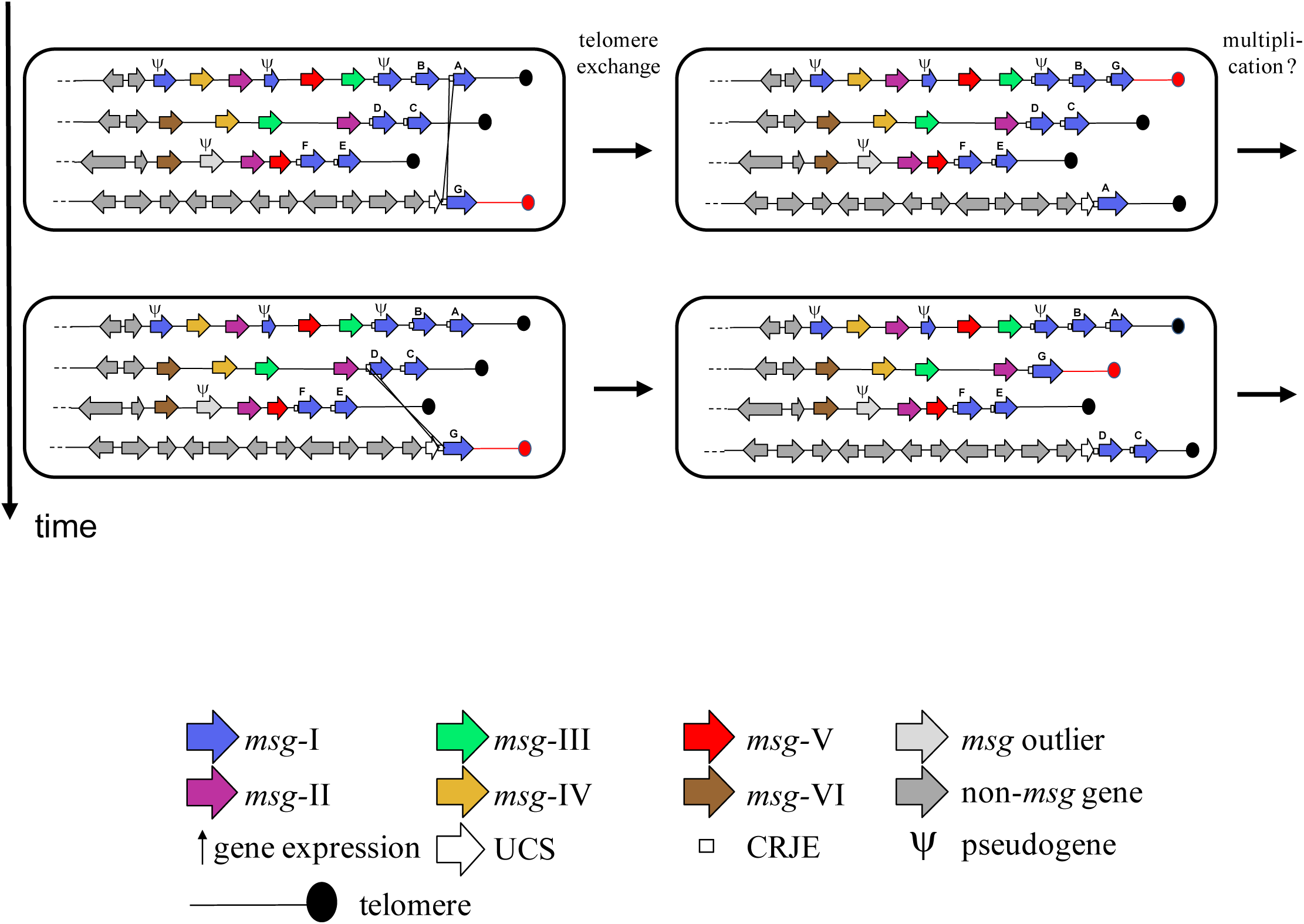
Telomere exchange model for swapping the *msg*-I expressed gene through a single recombination between CRJE sequences. One exchanged telomere is shown in red. Subpopulations of cells expressing a potentially new mosaic *msg*-I gene are generated over time and may then multiply. Polycistronic expression of two *msg*-I genes is figured in the second subpopulation generated (see text).

### Expression of *msg* families

RNAseq analyses suggested that the vast majority of the *msg* genes of all families were expressed in *P. carinii* and *P. murina* populations (4). As far as *P. jirovecii* is concerned, alignment of our previous RNAseq data (6) with the subtelomeres assembled in the present study was compatible with the same conclusion, although the data were from different clinical isolates (results not shown). Expression of most *msg*-I genes at the population level is consistent with the numerous subpopulations of cells expressing different *msg*-I genes that we observed. As far as *msg* families II to VI are concerned, the RNAseq data are compatible with constitutive or temporally regulated expression of all genes in each cell driven by the promoter present upstream of each of these genes. However, they are also compatible with mutually exclusive or partially exclusive expression of these genes thanks to silencing of promoters, or through another unknown mechanism.

### Cell surface structure

The UCS is a strong promoter (13), probably leading to a majority of a single isoform of adhesive Msg-I antigens on the cellular surface. This is consistent with the fact that Msg-I proteins are the most abundant at the cell surface (13). The surface of *P. carinii* trophic cells was shown to harbour also the surface protein INT1 participating to adhesion (30). Recently, a transcription factor responsible for expression of (a) still unidentified adhesive surface protein(s) has been reported in *P. carinii* trophic cells (31). Genes encoding orthologs of these two latter proteins are also present in the *P. jirovecii* genome (results not shown). Moreover, Kottom and Limper (31) mentioned that other uncharacterized genes which are important in binding to mammalian hosts are present in the *P. carinii* genome. Thus, the structure of *P. jirovecii* cell surface is made of a complex mixture of different proteins.

### Strategy of antigenic variation

The exchange of the *msg*-I isoform expressed and the generation of new mosaic genes of all *msg* families probably leads to a continuous segregation of subpopulations with a new mixture of glycoproteins at the cell surface. Thus, the strategy of the fungus would consist in the continuous generation of cells which are antigenically different. This strategy is further suggested by other characteristics of *Pneumocystis* spp. First, there is a high variability of the subtelomeres between *P. jirovecii* isolates (4) which is consistent with frequent subtelomeric recombinations. The subtelomeres of the isolate we studied here also differed greatly from those of the same chromosomes reported by Ma *et al* (4) (Supplementary note 6). Second, sexuality could be obligatory in the cell cycle (2, 3) because ectopic recombinations between subtelomeres occur during meiosis, within the bouquet of telomeres formed (8). The likely primary homothallic sexuality of *Pneumocystis* spp (32) avoids the need to find a compatible partner and thus increases mating frequency, which is believed to favor genetic diversity (33). Moreover, the genetic diversity might be enhanced by mating between the numerous co-infecting strains which are generally present in *P. jirovecii* infections (15). Third, the presence of several *msg* families may allow the formation of Msg hetero-oligomers that we envisage above, which could further enhance the cell surface complexity.

### Strategies of antigenic variation in different human pathogens

The mechanisms and hypothesized strategy of antigenic variation unravelled here appear unique among human pathogens. *Candida glabrata* contain one subtelomeric family of ca. 20 adhesins (7). *Trypanosoma brucei* presents a large reservoir of sequences used to create mosaic genes of a single surface antigen family made of about a thousand of genes located in subtelomeres as well as on minichromosomes (7). In the latter organism, pseudogenes provide segments to mosaic functional antigens (34), a phenomenon which might also occur in *P. jirovecii*. *Plamsodium falciparum* harbours one subtelomeric antigen family of ca. 60 members (7). These three organisms present a single gene family subject to mutually exclusive expression involving silencing in several cases. Thus, their populations are homogenous antigenically but may vary over time when the expressed gene is exchanged. Such strategy might be imposed by sterile niches such as blood and urinary tract. This contrasts sharply with the putative strategy of antigenic variation of *P. jirovecii* consisting in the continuous production of a mixture of cells antigenically different. The latter strategy may be associated to the particular niche within lungs since this niche tolerates the presence of low abundant fungi as members of the natural lung microbiota. This strategy might allow presenting most cells as different organisms to the immune system and thus to be tolerated during colonization. A similar strategy might be used by *Candida albicans* living in non-sterile mucosal niches. Indeed, its unique adhesin family presents a high number of serine CUG codons which are ambiguously translated into serine or leucine, thus creating variability from individual genes (35).

*Trypanosoma* and *Plasmodium* also differ from *Pneumocystis* spp in that they infect two different hosts rather than one. This undoubtedly exerts a different selective pressure on their antigenic variation system. The *Pneumocystis* spp differ considerably in their *msg* families (4), as well as in the fine structure of the Msg adhesins (36). It is likely that these differences are involved in the strict host species specificity of these fungi. Further work aiming at understanding the relation between structure and function of the different Msg glycoproteins is needed to further decipher both antigenic variation and host specificity of these fungi.

## Materials and Methods

### Ethics approval and consent to participate

The protocol was approved by the institutional review board (Commission cantonale d'éthique de la recherche sur l'être humain). All patients provided an informed written consent which was part of the procedure for admittance in the hospital. The admittance paperwork included the possibility to ask that their samples were not used for research. The samples were treated anonymously and were collected through a routine procedure at the hospital.

### Availability of data and materials

PacBio raw reads (accession number SRR5533719) and PacBio assembly (accession number pending) have been deposited in the NCBI Sequence Read Archive linked to BioProject PRJNA382815 and BioSample SAMN06733346. The datasets generated and analyzed during the current study are available from the corresponding author on reasonable request.

#### Bronchoalveolar lavage fluid specimens

Fresh BALFs positive for *P. jirovecii* using Methenamine-silver nitrate staining (37) were supplemented with 15% v/v glycerol, frozen in liquid Nitrogen, and stored at −80°C. Only those with more than one ml available and heavy fungal load were stored. Seventeen specimens were stored between 2012 and 2014, and used for the selection procedure described here below.

#### DNA extraction and identification of an infection with a single *P. jirovecii* strain

Genomic DNA was extracted from 0.2 to 0.4 ml of BALF specimen using QIAamp^®^ DNA Mini kit (Qiagen), and resuspended in 50 µl of elution buffer. Four genomic regions were amplified by PCR from genomic DNA extracted as described previously (38). Each PCR product was cloned into the plasmid pCR^tm^4-TOPO using the TOPO TA cloning Kit for Sequencing (Life Technologies). Both strands of the insert of 15 clones for each genomic region were sequenced with M13 primers using the BigDye Terminator kit and the ABI Prism 3100 automated sequencer (both from PerkinElmer Biosystems). Among the 17 clinical specimens collected, only one generated identical sequences for all clones of all genomic regions. Since ca. 15 clones per genomic region were analyzed, a second eventual co-infecting strain in this specimen should not represent more than ca. 7% of the *P. jirovecii* population. This specimen was selected for all experiments performed in the present study. It was from a HIV-infected patient.

#### Enrichment in *P. jirovecii* DNA and random amplification

The DNA of the selected specimen was enriched in *P. jirovecii* DNA using the NEBNext® Microbiome DNA Enrichment Kit based on the absence of CpG methylation (Biolabs), purified by ethanol precipitation in presence of 10 µg glycogen (Thermo Fisher Scientific), and resuspended in 50 µl of 1X TE Buffer. This enrichment raised the proportion of *P. jirovecii* DNA from a few percent to ca. 55% as determined *a posteriori* by high throughput sequencing. Because only small amounts of DNA are recoverable from a clinical specimen and in absence of an *in vitro* culture system, sufficient amount of DNA for high throughput PacBio sequencing was obtained by random amplification. Five µl of DNA was randomly amplified in a 50 µl reaction using the Illustra GenomiPhi HY DNA Amplification Kit (GE Healthcare). This amplification proved to create artificial molecules made of inverted repeats of several kb which were revealed by PacBio sequencing. The reads from these molecules were eliminated by bioinformatics (see below). DNA was then purified using QIAamp® DNA blood mini kit (Qiagen) followed by ethanol precipitation in presence of 10 µg glycogen. Amplified DNA fragments were sized (mean 8.6 kb) and quantified using Fragment Analyzer^TM^ (Advanced Analytical).

#### High throughput PacBio sequencing

Five µg of amplified DNA were used to prepare a SMRTbell library with the PacBio SMRTbell Template Prep Kit 1 according to the manufacturer's recommendations (Pacific Biosciences). The resulting library was size selected on a BluePippin system (Sage Science) for molecules larger than 5 kb. The recovered library was sequenced on one SMRT cell with P6/C4 chemistry and MagBeads on a PacBio RSII system (Pacific Biosciences) at 240 min movie length.

#### Read filtering and *P. jirovecii* genome assembly

The flow chart of the filtering and assembly procedure is shown in Figure S13a and the details for each step are described here. PacBio sub-reads were extracted from the raw h5-files using DEXTRACTOR (https://github.com/thegenemyers/DEXTRACTOR/). The average length of the extracted sub-reads was 5.2 kb with a maximum of 42 kb. We removed human derived reads by mapping them against the human reference genome using blasr (smrtpipe2.3, cut-off: corrected score < 55000). Reverse-complementary artificial reads created by the random amplification were next filtered out (cut-off:match length >=1000 bps) after mapping them onto themselves using DALIGNER (https://github.com/thegenemyers/DALIGNER/)(V1.0, options:-A -I). The cleaned reads were assembled using the tool FALCON (39) (V0.2, options: length_cutoff=8000m length_cutoff_pr=1000). PacBio reads were re-mapped onto the assembly using BLASR and used to evaluate and flag remaining human contigs. Human derived contigs were subsequently removed. A total of 2.2 Gb of *P. jirovecii* DNA sequences corresponding to a 200-fold coverage of the genome were gathered. The assembly was polished to remove residual PacBio errors using Quiver (40) (smrtpipe2.3, 5 iterations). The final polished genome assembly included 8.1 Mb in 219 gap-free contigs ranging from 234 bps to 386 kb with a NG50 of 108Kb, and 57% of the genome in 28 contigs lager than 100 kb. The *P. jirovecii* PacBio assembly obtained in the present study covered 96% of that we previously obtained using other sequencing methods^6^, and contained ca. 0.5 Mbp of subtelomeric sequences. The combination of both our assemblies covered 97% the assembly of Ma *et al* (4). Controls consisting in PCR amplification of specific subtelomeric regions from the same DNA sample confirmed the accuracy of the nucleotide sequence of the polished PacBio assembly, although few errors in repetitive homopolymer regions were detected (Supplementary note 8).

#### Gene predictions and *msg* annotations

Genes were predicted on the assembly using Augustus (41) (version 2.5.5) and a specifically trained model for *Pneumocystis* (6). In order to detect novel and more distant homologous *msg* genes in the assembly, we chose a generalized profile based approach (16) (Fig. S13b). A DNA profile was generated based on a previously described *msg* gene in *P. carinii* (42) (GenBank D82031.1) and a protein profile based on Msg-Rucl 21 (European Nucleotide Archive ABQ51002.1) using a Smith-Waterman-Algorithm (43). The profiles were calibrated against the scrambled genome (window approach, size=60). Using pfsearchV3 (44), the assembled genome was searched for homologues matches with the DNA profile. Curated matches were extracted and aligned against each other using MAFFT (45) (version 7.305). After manual curation and trimming, the alignments were divided in five groups based on neighbourhood joining (% identity) using Jalview (46) (v2.8.1). One representative candidate per group was selected and a new profile based on its sequence generated and calibrated as described here above. These DNA *msg* profiles were used to find and annotate a first set of 75 *msg* genes in the assembly. A combination of Blastx, genewise, in-house tools, and manual curation was applied using the protein Msg profile to extend and correct these annotations to the set of 113 *msg* genes analysed in the present study. The *msg* genes reported here were all manually curated with respect to their start, stop and intron coordinates.

#### Construction of phylogenetic trees

For the DNA and protein based phylogenetic analysis, the CDS for each annotated *msg* gene was manually corrected (up to five corrections), extracted, and translated into its protein sequence. Both CDS and protein sequences were aligned against each other using MAFFT (45) (mafft-linsi –genafpair), and the multiple sequence alignment used to infer a phylogenetic tree with RAxML (47) (PROTGAMMAGTR for proteins and with GTRGAMMA for CDS, 1000 bootstraps). The *msg* genes of family V were defined as out-group and the final tree rooted. Proteins were further classified using JACOP (48) (http://myhits.isb-sib.ch/cgi-bin/jacop/). In order to add pseudogenes and published *msg* genes from Ma *et al* (4) equal or exceeding 1.6 kb, we injected the new sequences into the prior DNA-based multiple-alignment using MAFFT (45) (--addfull). They were added to the original tree using the evolutionary placement algorithm (EPA) from RAxML. These trees were converted into a compatible format with the tool guppy from the pplacer suite (49) (v1.1alpha14, tog). Genes with an exon smaller than 1.6 kb were added to the original DNA-based multiple-alignment using MAFFT (45) (--addfragments). The alignment was trimmed and re-aligned using MAFFT (45). A new tree was then build with RAxML (GTRGAMMA, 1000 bootstraps). All trees were analyzed and visualized using R (50) (3.3.2) and GGTREE (51) (v1.6.9).

#### Gene and protein sequences analyses

Alignments of full-length gene or protein sequences were carried out using MAFFT (43). Canonical TATA box and Cap signal (52), as well as canonical donor and acceptor sequences of *Pneumocystis* introns (53, 54), were identified by visual inspection of the alignments and sequences of the genes. Signal peptide and GPI anchor signal were identified using respectively Phobius (55) (http://phobius.sbc.su.se/) and GPI-SOM (56) (http://gpi.unibe.ch/) with default settings. Canonical potential sites NXS/T of Nitrogen-linked glycosylation (21) were identified by visual inspection. Conserved domains were searched using Multiple Expectation–Maximization for Motif Elicitation (17) (MEME, http://meme-suite.org/tools/meme). MEME analysis of the 49 full-length Msg proteins of all families except outliers was carried out using default settings, except minimum and maximum motif width of respectively 50 and 100 residues, any number of sites per sequence option, and maximum of 13 motifs searched. HMMER (57) (biosequence analysis using profile hidden Markov models, http://www.ebi.ac.uk/Tools/hmmer/search/hmmscan) was used with default settings on full-length proteins for the following embedded predictions: Pfam, unstructured regions (Intrinsically Unstructured Proteins, IUPRED), and coiled-coil motifs (Ncoils predictor). Pairwise identities between full-length *msg* genes and Msg proteins were calculated using the multi-way alignment type of Clone Manager 9 professional edition software.

#### Search for potential mosaic genes

Two screening methods were first used: Recombination Analysis tool (58) (RAT, http://cbr.jic.ac.uk/dicks/software/RAT/) and Bellerophon (59) (http://comp-bio.anu.edu.au/bellerophon/bellerophon.pl). MAFFT (45) alignments of various set of genes were analysed with both methods. RAT was used with default settings, *i.e.* using windows of one tenth of the length of the alignment and increment size equal to half of the window size. Bellerophon was used with default settings, *i.e.* windows of 300 bps and Huber-Hugenholtz correction. RAT can detect several recombination events whereas Bellerophon reports a single one per mosaic gene. The more sensitive method TOPALi v2.5 (60) (http://www.topali.org/) which is based on a Hidden Markov Model (HMM) was then applied on the potential mosaic genes and its putative parent genes detected with the two screening methods. These three genes were aligned using MAFFT (45) with an additional gene chosen randomly in the same *msg* family since TOPALi requires four genes in input. The efficacy of the three methods to detect mosaic genes was assessed by the analysis of artificial chimera produced *in silico* with related genes, as well as with sets of orthologous genes from different fungal species (results not shown). Only the RAT method is suitable for the search of recombination events among proteins. The vast majority of the events detected at the protein level corresponded to those detected at the DNA level (results not shown).

#### PCR amplification and sequencing

PCRs were performed in a final volume of 20 µl with 0.35 U of High Fidelity Expand polymerase (Roche Diagnostics), using the buffer provided, each dNTP at a final concentration of 200 µM, and each primer at 0.4 µM. PCR conditions included an initial denaturation step of 3 min at 94°C, followed by 35 cycles consisting of 30 s at 94°C, 30 s at the annealing temperature, and 1 min per kb to be amplified at 72°C. The reaction ended with 5 min of extension at 72°C. The annealing temperature and the MgCl_2_ concentration were optimized for each set of primers and ranged from 51 to 60°C and from 3 to 6 mM, respectively. Sequencing both strands of the PCR products was performed with the two primers used for PCR amplification, as well as the Big Dye Terminator DNA sequencing kit and ABI PRISM 3100 automated sequencer (both from Perkin-Elmer Biosystems).

#### Accessibility of data and materials

PacBio raw reads (accession number SRR5533719) and PacBio assembly (NJFV00000000) have been deposited in the NCBI Sequence Read Archive linked to BioProject PRJNA382815 and BioSample SAMN06733346. The datasets generated and analyzed during the current study are available from the corresponding author on reasonable request.

#### Funding information

This work was supported by the Swiss National Science Foundation, grant 310030_146135 to P.M.H. and M.P. This Foundation had not role in any steps of the study.

## Acknowledgments

Computations were performed at the Vital-IT Center for High-Performance Computing of the Swiss Institute of Bioinformatics (http://www.vital-it.ch). We thank Michel Monod, Dominique Sanglard, and Laurent Keller for critical reading.

## References

1. Cushion MT, Smulian AG, Slaven BE, Sesterhenn T, Arnold J, Staben C, Porollo A, Adamczak R, Meller J. 2007. Transcriptome of *Pneumocystis carinii* during fulminate infection: carbohydrate metabolism and the concept of a compatible parasite. PLoS ONE 2: e423.

2. Cushion MT, Stringer JR. 2010. Stealth and opportunism: alternative lifestyles of species in the fungal genus *Pneumocystis*. Annu Rev Microbiol 64: 431–452.

3. Hauser PM. 2014. Genomic insights into the fungal pathogens of the genus *Pneumocystis*: obligate biotrophs of humans and other mammals. PLoS Pathog 10: e1004425.

4. Ma L, Chen Z, Wei Huang D, Kutty G, Ishihara M, Wang H, Abouelleil A, Bishop L, Davey E, Deng R, Deng X, Fan L, Fantoni G, Fitzgerald M, Gogineni E, Goldberg JM, Handley G, Hu X, Huber C, Jiao X, Jones K, Levin JZ, Liu Y, Macdonald P, Melnikov A, Raley C, Sassi M, Sherman BT, Song X, Sykes S, Tran B, Walsh L, Xia Y, Yang J, Young S, Zeng Q, Zheng X, Stephens R, Nusbaum C, Birren BW, Azadi P, Lempicki RA, Cuomo CA, Kovacs JA. 2016. Genome analysis of three *Pneumocystis* species reveals adaptation mechanisms to life exclusively in mammalian hosts. Nat com 7: 10740.

5. Brown GD, Denning DW, Gow NAR, Levitz SM, Netea MG, White TC. 2012. Hidden killers: human fungal infections. Sci Transl Med 4: 165rv13.

6. Cissé OH, Pagni M, Hauser PM. 2012. *De novo* assembly of the *Pneumocystis jirovecii* genome from a single bronchoalveolar lavage fluid specimen from a patient. mBio 4: e00428–12.

7. Deitsch KW, Lukehart SA, Stringer JR. 2009. Common strategies for antigenic variation by bacterial, fungal and protozoan pathogens. Nat Rev 7: 493–503.

8. Barry JD, Ginger ML, Burton P, McCulloch R. 2003. Why are parasite contingency genes often associated with telomeres? Int. J. Parasit. 33: 29–45.

9. Keely SP, Renauld H, Wakefield AE, Cushion MT, Smulian AG, Fosker N, Fraser A, Harris D, Murphy L, Price C, Quail MA, Seeger K, Sharp S, Tindal CJ, Warren T, Zuiderwijk E, Barrell BG, Stringer JR, Hall N. 2005. Gene arrays at *Pneumocystis carinii* telomeres. Genetics 170: 1589–1600.

10. Keely SP, Stringer, JR. 2009. Complexity of the MSG gene family of *Pneumocystis carinii*. BMC Gen 10: 367.

11. Stringer JR. 2007. Antigenic Variation in *Pneumocystis*. J Eukaryot Microbiol 54: 8–13.

12. Kutty G, Ma L, Kovacs JA. 2001. Characterization of the expression site of the major surface glycoprotein of human-derived *Pneumocystis carinii*. Mol Microbiol 42: 183–193.

13. Kutty G, Shroff R, Kovacs JA. 2013. Characterization of *Pneumocystis* major surface glycoprotein gene (*msg*) promoter activity in *Saccharomyces cerevisiae*. Euk Cell 12: 1349–1355.

14. Kutty G, Maldarelli F, Achaz G, Kovacs JA. 2008. Variation in the major surface glycoprotein genes in *Pneumocystis jirovecii*. J Infect Dis 198: 741–749.

15. Alanio A, Gits-Muselli M, Mercier-Delarue S, Dromer F, Bretagne S. 2016. Diversity of *Pneumocystis jirovecii* during infection revealed by ultra-deep pyrosequencing. Front Microbiol 7: 733.

16. Bucher P, Bairoc A. 1994. A generalized profile syntax for biomolecular sequence motifs and its function in automatic sequence interpretation. Proc Int Conf Intell Syst Mol Biol 2: 53–61.

17. Bailey TL, Elkan C. 1994. Fitting a mixture model by expectation maximization to discover motifs in biopolymers. Proc Sec Int Conf Int Syst Mol Biol. 28–36 (AAAI Press, Menlo Park, California).

18. Pottratz ST, Paulsrud J, Smith JS, Martin WJ II. 1991. *Pneumocystis carinii* attachment to cultured lung cells by *Pneumocystis* gp 120, a fibronectin binding protein. J Clin Invest 88: 403–407.

19. Limper AH, Standing JE, Hoffman OA, Castro M, Neese, LW. 1993. Vitronectin binds to *Pneumocystis carinii* and mediates organism attachment to cultured lung epithelial cells. Infect Immun 61: 4302–4309.

20. Dranginis AM, Rauceo JM, Coronado JE, Lipke PN. 2007. A Biochemical Guide to Yeast Adhesins: Glycoproteins for Social and Antisocial Occasions. Microbiol Mol Biol Rev 71: 282–294.

21. Linder T, Gustafsson CM. 2008. Molecular phylogenetics of ascomycotal adhesins—a novel family of putative cell-surface adhesive proteins in fission yeasts. Fung Gen Biol 45: 485–497.

22. Hakoshima T. Leucine Zippers. 2005. eLS.

23. Williamson MP. 1994. The structure and function of proline-rich regions in proteins. Biochem J 297: 249–260.

24. Strauss HM, Keller S. 2008. Pharmacological interference with protein-protein interactions mediated by coiled-coil motifs. Hand. Exp Pharmacol 186: 461–482.

25. Hitchcock-DeGregori SE, Barua B. 2017. Tropomyosin structure, function, and interactions: a dynamic regulator. Sub Biochem 82: 253–264.

26. Best RB. 2017. Computational and theoretical advances in studies of intrinsically disordered proteins. Curr Opin Struct Biol 42:147–154.

27. Sunkin SM, Stringer JR. 1996. Translocation of surface antigen genes to a unique telomeric expression site in *Pneumocystis carinii*. Mol Microbiol 19: 283–295.

28. Hua SB, Qiu M, Chan E, Zhu L, Luo Y. 1997. Minimum length of sequence homology required for *in vivo* cloning by homologous recombination in yeast. Plasmid 38: 91–96.

29. Turan S, Bode J. 2011. Site-specific recombinases: from tag-and-target-to tag-and-exchange-based genomic modifications. FASEB J 25: 4088–4107.

30. Kottom TJ1, Kennedy CC, Limper AH. 2008. *Pneumocystis PCINT1*, a molecule with integrin-like features that mediates organism adhesion to fibronectin. Mol Microbiol 67: 747–761.

31. Kottom TJ, Limper AH. 2016. Evidence for a *Pneumocystis carinii* Flo8-like transcription factor: insights into organism adhesion. Med Microbiol Immunol 205: 73–84.

32. Almeida JMGCF, Cissé OH, Fonseca Á, Pagni M, Hauser PM. 2015. Comparative genomics suggests primary homothallism of *Pneumocystis* species. mBio 6: e02250–14.

33. Roach KC, Heitman J. 2014. Unisexual reproduction reverses Muller’s ratchet. Genetics 198: 1059–1069.

34. Hall JPJ, Wang H, Barry JD. 2013. Mosaic *VSGs* and the scale of *Trypanosoma brucei* antigenic variation. PLoS Pathog 9: e1003502.

35. Rizzetto L, Weil T, Cavalieri D. 2015. Systems level dissection of *Candida* recognition by dectins: a matter of fungal morphology and site of infection. Pathog 4: 639–661.

36. Mei Q, Turner RE, Sorial V, Klivington D, Angus CW, Kovacs JA. 1998. Characterization of major surface glycoprotein genes of human *Pneumocystis carinii* and high-level expression of a conserved region. Infect Immun 66: 4268–4273.

37. Musto L, Flanigan M, Elbadawi A. 1982. Ten-minute silver stain for *Pneumocystis carinii* and fungi in tissue sections. Arch Pathol Lab Med 106: 292–294.

38. Hauser PM, Francioli P, Bille J, Telenti A, Blanc DS. 1997. Typing of *Pneumocystis carinii* f. sp. *hominis* by single-strand conformation polymorphism of four genomic regions. J Clin Microbiol 35: 3086–3091.

39. Chin CS, Peluso P, Sedlazeck FJ, Nattestad M, Concepcion GT, Clum A, Clum A, Dunn C, O’Malley R, Figueroa-Balderas R, Morales-Cruz A, Cramer GR, Delledonne M, Luo C, Ecker JR, Cantu D, Rank DR, Schatz MC. 2016. Phased diploid genome assembly with single-molecule real-time sequencing. Nat Meth 13: 1050–1054.

40. Chin CS, Alexander DH, Marks P, Klammer AA, Drake J, Heiner C, Clum A, Copeland A, Huddleston J, Eichler EE, Turner SW, Korlach J. 2013. Nonhybrid, finished microbial genome assemblies from long-read SMRT sequencing data. Nat Meth 10: 563–569.

41. Stanke M, Keller O, Gundunz I, Hayes A, Waack S, Morgenstern B. 2006. AUGUSTUS: *ab initio* prediction of alternative transcripts. Nucl Ac Res 34: W435–439.

42. Wada M, Nakamura Y. 1996. Unique telomeric expression site of major-surface-glycoprotein genes of *Pneumocystis carinii*. DNA Res 3: 55–64.

43. Smith TF, Waterman MS. 1981. Identification of common molecular subsequencees. J Mol Biol 147: 195–197.

44. Schuepbach T, Pagni M, Bridge A, Bouqueleret L, Xenarios I, Cerutti L. 2013. pfsearchV3: a code acceleration and heuristic to search PROSITE profiles. Bioinfo 29: 1215–1217.

45. Katoh K, Standley DM. 2013. MAFFT multiple sequence alignment software version 7: improvements in performance and usability. Mol Biol Evol 30: 772–780.

46. Waterhouse AM, Procter JB, Martin DM, Clamp M, Barton GJ. 2009. Jalview version 2-a multiple sequence alignment and analysis workbench. Bioinfo 25: 1189–1191.

47. Stamatakis A. 2014. RAxML version 8: a tool for phylogenetic analysis and post-analysis of large phylogenies. Bioinfo 30: 1312–3.

48. Sperisen P, Pagni M. 2005. JACOP: a simple and robust method for the automated classification of protein sequences with modular architecture. BMC Bioinfo 6: 216.

49. Matsen FA, Kodner RB, Armbrust, EV. 2010. pplacer: linear time maximum-likelihood and Bayesian phylogenetic placement of sequences onto a fixed reference tree. BMC Bioinfo 11: 538.

50. R Core Team. 2013. R: a language and environment for statistical computing. R Foundation for Statistical Computing, Vienna, Austria. URL http://www.R-project.org/.

51. Yu G, Smith DK, Zhu H, Guan Y, Lam TT-Y. 2017. GGTREE: an R package for visualization and annotation of phylogenetic trees with their covariates and other associated data. Meth Ecol Evol 8: 28–36.

52. Bucher P. 1990. Weight matrix descriptions of four eukaryotic RNA polymerase II promoter elements derived from 502 unrelated promoter sequences. J Mol Biol 212: 563–578.

53. Thomas CF JR., Loef EB, Limper AH. 1999. Analysis of *Pneumocystis carinii* introns. Infect Immun 67: 6157–6160.

54. Slaven BE, Porollo A, Sesterhenn T, Smulian AG, Cushion MT, Meller J. 2006. Large-scale characterization of introns in the *Pneumocystis carinii* genome. J Eukar Microbiol 53: S151–153.

55. Käll L, Krogh A, Sonnhammer ELL. 2004. A Combined Transmembrane Topology and Signal Peptide Prediction Method. J Mol Biol 338: 1027–1036.

56. Fankhauser N, Mäser P. 2005. Identification of GPI anchor attachment signals by a Kohonen self-organizing map. Bioinfo 21: 1846–1852.

57. Finn RD, Clements J, Eddy SR. 2011. HMMER web server: interactive sequence similarity searching. Nucl Ac Res 39: W29–37.

58. Etherington GJ, Dicks J, Roberts IN. 2005. Recombination Analysis Tool (RAT): a program for the high-throughput detection of recombination. Bioinfo 21: 278–281.

59. Huber T, Faulkner G, Hugenholtz P. 2004. Bellerophon: a program to detect chimeric sequences in multiple sequence alignments. Bioinfo 20: 2317–2319.

60. Milne I, Wright F, Rowe G, Marshal DF., Husmeier D, McGuire G. 2004. TOPALi: software for automatic identification of recombinant sequences within DNA multiple alignments. Bioinfo 20: 1806–1807.

61. Ma L, Kutty G, Jia Q, Imamichi H, Huang L, Atzori C, Beckers P, Groner G, Beard CB, Kovacs JA. 2002. Analysis of variation in tandem repeats in the intron of the major surface glycoprotein expression site of the human form of *Pneumocystis carinii*. J Infect Dis 186: 1547–1554.

62. Ramana J, Gupta, D. 2010. FaaPred: A SVM-based prediction method for fungal adhesins and adhesin-like proteins. PLoS ONE 5:e9695.

